# Evaluation of Antibiotic Tolerance in *Pseudomonas aeruginosa* for Aminoglycosides and its Predicted Gene Regulations Through In-silico Transcriptomic Analysis

**DOI:** 10.1101/2021.05.01.442230

**Authors:** Abishek Kumar, Bency Thankappan, Angayarkanni Jayaraman, Akshita Gupta

## Abstract

*Pseudomonas aeruginosa* causes chronic infections like cystic fibrosis, endocarditis, bacteremia and sepsis, which are life-threatening and difficult to treat. The lack of antibiotic response in *P. aeruginosa* is due to adaptive resistance mechanism, which prevents the entry of antibiotics into cytosol of the cell to achieve tolerance. Among different groups of antibiotics, aminoglycosides are used as a parental antibiotic for treatment of *P. aeruginosa*. This study aims to determine the kinetics of antibiotic tolerance and gene expression changes in *P. aeruginosa* exposed to amikacin, gentamicin, and tobramycin. These antibiotics were exposed to *P. aeruginosa* at their MICs and the experimental setup was monitored till 72 hours, followed by the measurement of optical density in the interval of every 12 hours. The growth of *P. aeruginosa* in MICs of antibiotics represents the kinetics of antibiotic tolerance in amikacin, gentamicin, and tobramycin. Transcriptomic profile of antibiotic exposed *P. aeruginosa* PA14 was taken from Gene Expression Omnibus (GEO), NCBI as microarray datasets. The gene expressions of two datasets were compared by test versus control. Tobramycin exposed *P. aeruginosa* failed to develop tolerance in MICs 0.5µg/mL, 1µg/mL and 1.5µg/mL. Whereas amikacin and gentamicin treated *P. aeruginosa* developed tolerance in MICs. This depicts the superior *in vitro* response of tobramycin over the gentamicin and amikacin. Further, *in silico* transcriptomic analysis of tobramycin treated *P. aeruginosa* resulted in low expression of 16s rRNA Methyltransferase E, B & L, alginate biosynthesis genes and several proteins of Type 2 Secretory System (T2SS) and Type 3 Secretory System (T3SS). The Differentially Expressed Genes (DEGs) of alginate biosynthesis, and RNA Methyltransferases suggests increased antibiotic response and low probability of developing resistance. The use of tobramycin as a parental antibiotic with its synergistic combination might combat *P. aeruginosa* with increased response.

## 1. Introduction

*P. aeruginosa* is an opportunistic pathogen, causes chronic infections which are difficult to treat because of the limited response to antimicrobials and emergence of antibiotic resistance during therapy[1]. Multi-drug resistance (MDR) in *P. aeruginosa* is increasing due to over-exposure to antibiotics[2]. It is developed by various physiological and genetic mechanisms, which includes multidrug efflux pumps, beta-lactamase production, outer membrane protein (porin) loss and target mutations. In hospitals, MRD *P. aeruginosa* are concurrently resistant to ciprofloxacin, imipenem, ceftazidime and piperacillin-tazobactam in most of the cases, which limits the treatment options[3].

Aminoglycosides are major group of antibiotics with potential bacteriocidic effect for the treatment of *Pseudomonas* infections. They are either used alone or in combination to treat various infections to overcome drug resistance, particularly in cystic fibrosis patients[4], and infective endocarditis[5]. In the face of systemic infection with shock/sepsis, antimicrobial therapy should consist of two antimicrobial agents, with one of these being an aminoglycoside[6], because it exhibits concentration-dependent bactericidal activity and produce prolonged post-antibiotic effects[7].

β-lactam antibiotic plus an aminoglycoside is the commonly used synergistic combinations for treatment of clinical infections. Other combinations are fluoroquinolone & aminoglycosides and tetracycline & aminoglycosides. Clinical isolates show high percent of susceptibility to aminoglycosides than the other first-line antibiotics[8]. Despite high susceptibility to aminoglycosides in clinical isolates, *P. aeruginosa* exhibits physiological adaptations to the antibiotics which results in less response to its synergistic combinations.

*P. aeruginosa* thrives in the inhibitory concentration of antibiotics gradually and acquires adaptive resistance, which makes the treatment more complicated[9]. Adaptive resistance mechanism was characterized by modification of the cytoplasmic membrane, condensation of membrane proteins and reduction of phospholipid content[10] which reduces penetration of antibiotics into the plasma membrane. Studies shows that the adaptive resistance can also develop by up-regulation of efflux pumps especially, MexXY-OprM[11].

Among immunocompromised patients, *P. aeruginosa* are favored to adapt the administered antibiotics and enable better survival of the bacterial generation by emerging as physiologically resistant groups[12]. Adaptive resistance is developed due to rapid transcriptomic alteration in response to antibiotic[13]. Better understanding of the kinetics and transcriptomic changes during antibiotic exposure can develop scientific insight on the adaptive resistance mechanism in *P. aeruginosa*[13,14]. On this background our study was designed for better understanding of adaptive resistance in *P. aeruginosa* for gentamicin, amikacin and tobramycin which are commonly used aminoglycosides as first line antibiotics.

## 2. Materials and Methods

### 2.1 Broth Dilution Method

Minimum Inhibitory Concentrations (MICs) of *P. aeruginosa* ATCC 27853 was determined by broth dilution method as per the Clinical and Laboratory Standards Institute (CLSI) guidelines. MIC assay was performed for gentamicin, amikacin, and tobramycin (purchased from Sigma Aldrich) with log phase culture (5×10^8^ CFU/mL) in Mueller Hinton Broth (MHB) using 96-well microtiter plate. The final optical density (OD) was determined in Epoch^™^ Microplate spectrophotometer at 600nm.

### 2.2 In vitro exposure of antibiotics to P. aeruginosa

From the recorded MIC values, *P. aeruginosa* was inoculated in 10 mL MHB in a Tarsons tube with corresponding antibiotic concentrations after adjusting the cell density to OD_600_ 0.26 (in log phase). The experiment set up was observed for 72 hours. *P. aeruginosa* was inoculated in antibiotic concentrations 0.5μg/mL, 1μg/mL & 1.5μg/mL for gentamicin & tobramycin. For amikacin, 1μg/mL, 2μg/mL & 3μg/mL antibiotic concentrations were taken. All antibiotic concentrations taken were based on MICs determined for *P. aeruginosa* ATCC 27853. The experimental condition was incubated at 37°C with optimal shaking of 74rpm. At every 12 hours the turbidity was monitored by measuring OD_600_ in Epoch^™^ microtiter plate reader[14][15]. The tube with growth were sub-cultured in nutrient agar. The colonies were conformed for *P. aeruginosa* by Matrix-assisted laser desorption/ionization time-of-flight (MALDI-TOF) automated identification system (VITEK® MS, BioMérieux) by following the standard procedure for sample preparation[16].

### 2.3 In silico transcriptomic analysis

#### 2.3.1 Retrieval of microarray datasets

The differential gene expression analysis was performed by exploring microarray datasets published in Gene Expression Omnibus (GEO) NCBI, available as accessible series no. GSE9991 and GSE9989. Two datasets 1.) Tobramycin treated planktonic culture of *P. aeruginosa* (GSE9991) and 2.) Tobramycin treated *P. aeruginosa* biofilm (GSE9989) were analyzed. In <u>GSE9991</u>, planktonic culture of PA14 was exposed to 5 µg/mL tobramycin for 30 minutes at 37^0^C.Two samples GSM252561 and GSM252562 of tobramycin treated PA14 planktonic culture was taken as test, which was compared to 2 samples GSM252559 and GSM252560 of unexposed PA14 planktonic culture as control. <u>GSE9989</u> consist of 6 samples, in which 3 samples GSM252496, GSM252501 and GSM252505 of unexposed *P. aeruginosa* biofilm were taken as control and 3 samples GSM252506, GSM252507 and GSM252508 of Tobramycin exposed *P. aeruginosa* biofilm was taken as test. Biofilms were grown on CFBE41o-cells in culture for 9 hours in MEM/0.4% arginine. Replicate samples were then incubated in the presence or absence of 500 μg/mL tobramycin for 30 minutes[17].

In the published datasets, before RNA harvesting, cells were washed several times in 2 ml PBS to remove antibiotics. The RNA was extracted using RNeasy RNA isolation kit. Bacterial RNA was then purified using MicrobEnrich kit, to exclude mammalian RNA. The purified RNA was subjected to cDNA synthesis followed by microarray preparation according to Affymetrix - Genechip *P. aeruginosa* Genome Array Expression Analysis Protocol[17].

#### 2.3.2 Differential gene expression analysis

In the microarray datasets, raw data available as platform file was extracted, formatted, and uploaded as input data in NetworkAnalyst 3.0 (https://www.networkanalyst.ca). In NetworkAnalyst 3.0, organism ID – *P. aeruginosa*, data type – intensity table (Microarray data) options were selected, and probe summarization was performed by multi-array average algorithm. The raw data was preprocessed by removing unannotated genes, and datasets were normalized by log2 transformation. By Linear Models for Microarray Analysis (Limma) statistical package, datasets were subjected to specific comparison by its test versus control samples to determine log2 fold change (LogFc) between two groups. To identify significant DEGs, the datasets were filtered by significant threshold cutoff of P-value ≤0.05[18].

#### 2.3.3 Gene Ontology (GO) and functional enrichment analysis

PANTHER Classification System (http://pantherdb.org/)[19] and DAVID Bioinformatics Resources 6.8 (https://david.ncifcrf.gov/)[20] computational tools were used for further downstream analysis of significant DEGs. Gene ontology was performed in PANTHER for functional classification of DEGs under three categories which includes, molecular functions, biological process, and protein class. For enrichment analysis of significant DEGs, DAVID Bioinformatics Resources 6.8 tool was used. The databases selected for enrichment analysis in DAVID were KEGG Pathway, InterPro, UniProtKB, and SMART. Minimum threshold gene counts of ≥ 2 and EASE Score of 0.1 were set as cut off to filter and enrichments of P-value ≤ 0.05 and false discovery rate (FDR) ≤ 0.05 were only considered for the study. In the functional annotation clusters, enrichment score of 2.0 were set as threshold to select enriched clusters.

Further, target genes were annotated functionally using Pseudomonas Genome DB (http://pseudomonas.com/)[21] and PseudoCyc (http://www.pseudomonas.com:1555)[22] to determine the protein and molecular interactions of DEGs based on previously published resources.

## 3. Results

### 3.1. Broth Dilution Method

MICs of gentamicin, amikacin and tobramycin for *P. aeruginosa* (ATCC 27853) were 0.5 μg/mL, 1.5 μg/mL and 0.5 μg/mL.

### 3.2. *In vitro* exposure of antibiotics to *P. aeruginosa*

The initial OD_600_ of the bacterial culture was _∼_ 0.26 in all the tubes and after 12 hours the OD_600_ dropped to ∼0.13. In 0.5 μg/mL & 1 μg/mL of gentamicin (Figure 1) and 1 μg/mL & 2 μg/mL of amikacin (Figure 3) tubes, OD_600_ increased exponentially after 24 hours. After 48 hours, the bacterial growth attained the initial OD_600_ ∼0.26 (Figures 1 and 3) and the cells resumed active growth after the post-antibiotic effect. In tobramycin and higher concentrations of gentamicin and amikacin tubes, after 12 hours OD_600_ further declined, and no growth were observed till 72 hours (Figures 1, 2 and 3). The growth in 1 μg/mL & 2 μg/mL of amikacin and 0.5 μg/mL & 1 μg/mL of gentamicin tubes were confirmed as *P. aeruginosa* by MALDI-TOF automated identification system, based on the peptide mass fingerprint matching.

**Figure 1:**
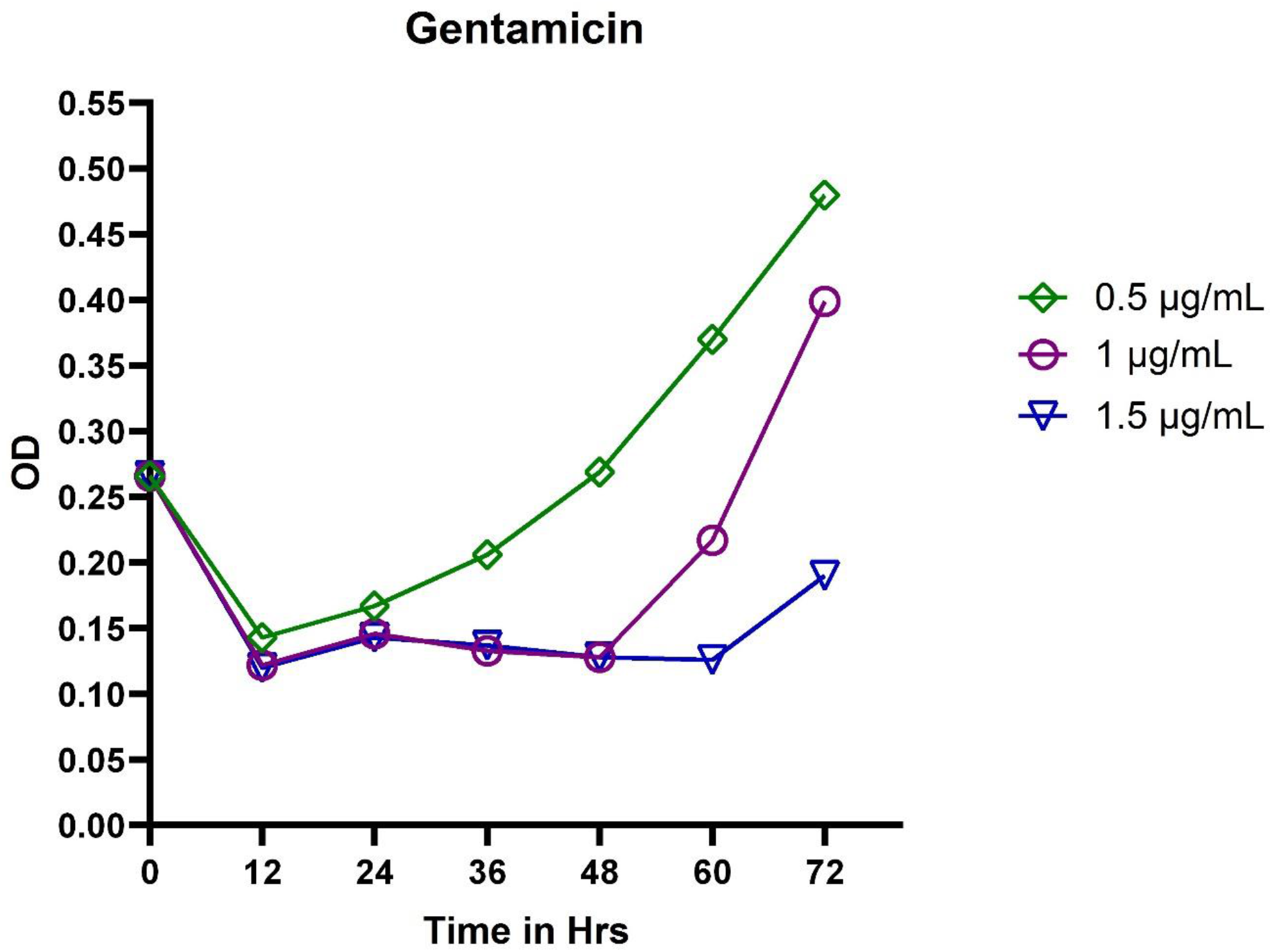
*In vitro* exposure of MICs 0.5μg/mL, 1μg/mL, and 1.5μg/mL of gentamicin to *P. aeruginosa*. The OD values depict the kinetics of adaptive resistance in the cell.

**Figure 2:**
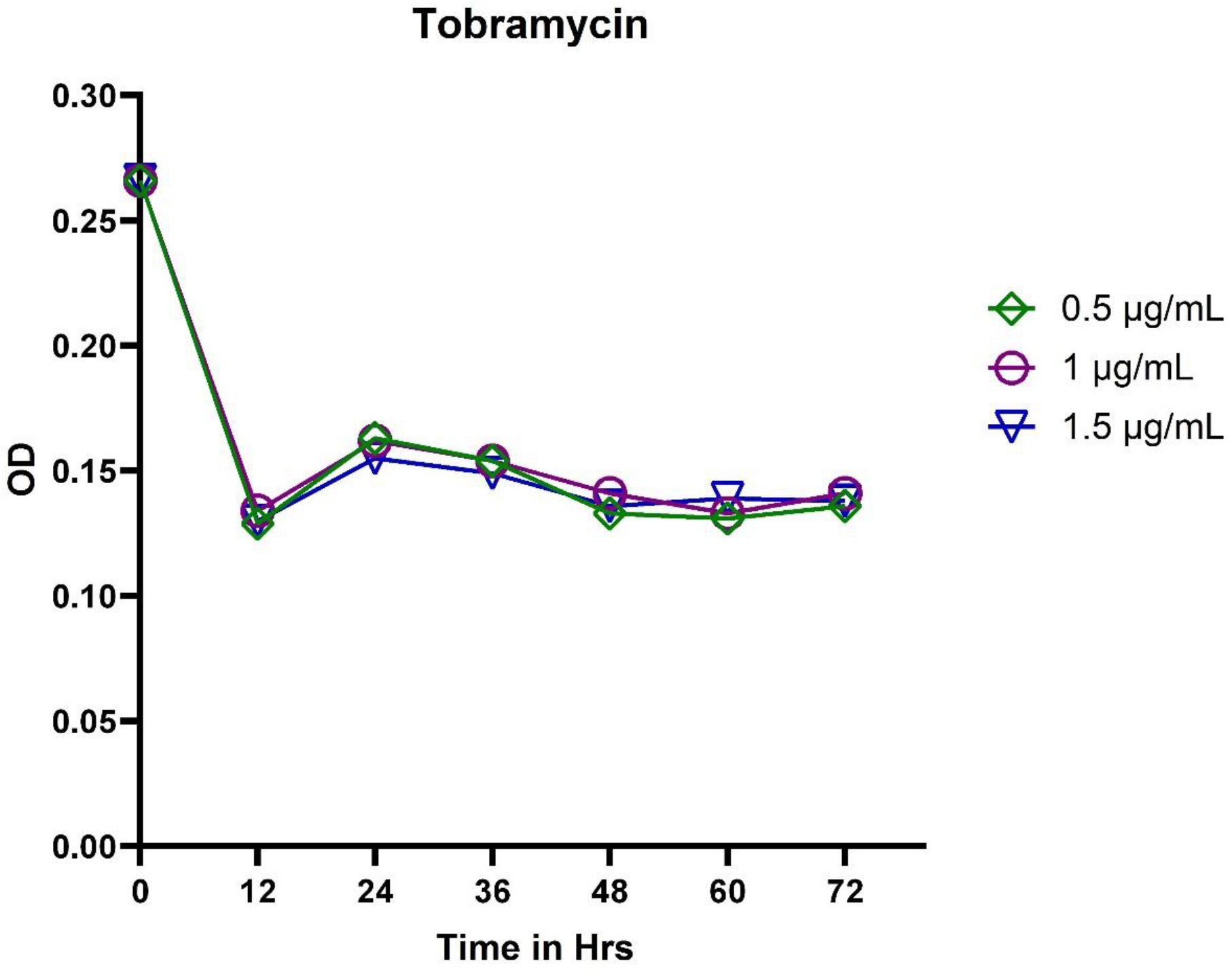
*In vitro* exposure of MICs 0.5 μg/mL, 1 μg/mL, and 1.5 μg/mL of tobramycin to *P. aeruginosa*. The OD values depict the kinetics of adaptive resistance in the cell.

**Figure 3:**
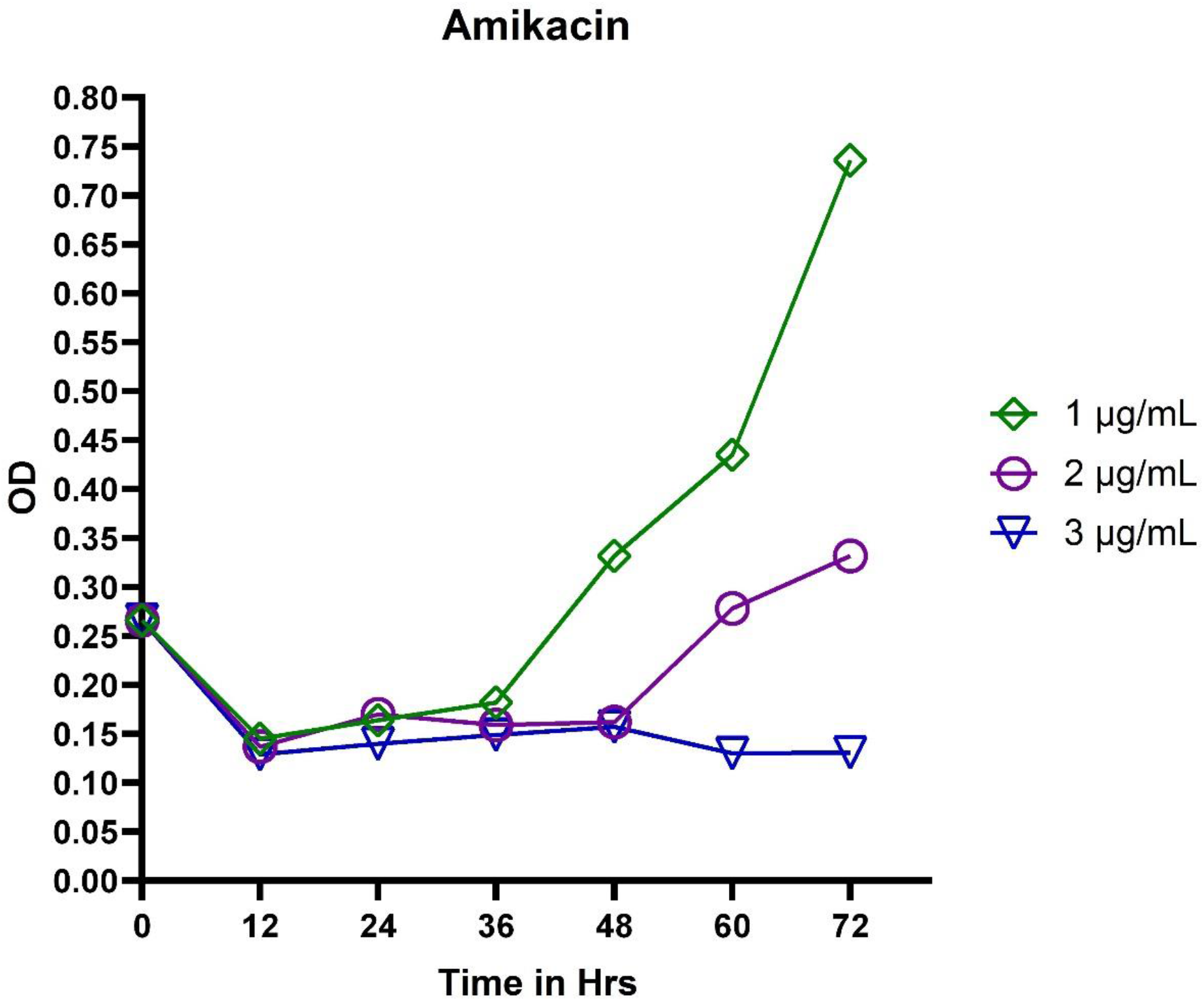
*In vitro* exposure of MICs 1 μg/mL, 2 μg/mL, and 3 μg/mL of amikacin to *P. aeruginosa*. The OD values depict the kinetics of adaptive resistance in the cell.

### 3.3. Differential gene expression analysis

In <u>GSE9991</u>, among 125 of DEGs 53 genes were upregulated and 72 were downregulated. In <u>GSE9989</u>, a total of 307 genes were differentially expressed in which 52 genes were upregulated and 255 genes were downregulated. Distribution of DEGs in both the datasets were represented in volcano plot (Figure 4). Targeted DEGs in the study were 17 from GSE9991 and 22 from GSE9989 (Table 1&2).

**Figure 4:**
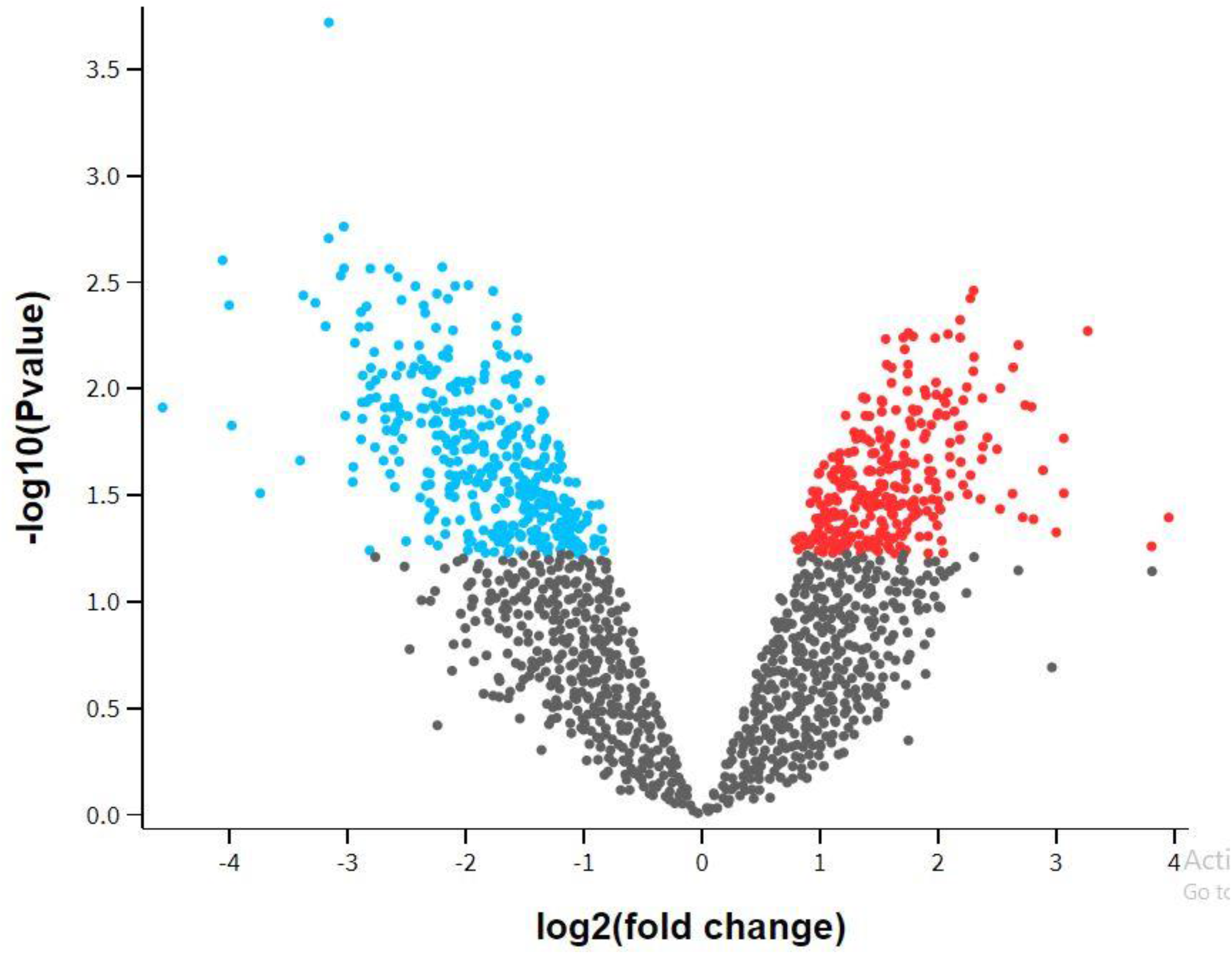
Volcano plot representing the distribution of DEGs in tobramycin treated *P. aeruginosa*. The figure represents the degree of variation in the gene expression of *P. aeruginosa* after exposure of tobramycin.

**Table 1.** List of target DEGs and its log fold change of expression in tobramycin treated planktonic *P. aeruginosa* (GSE9991)

| Genes | Log Fc |
| --- | --- |
| PA2677 | -2.21809 |
| PA0687 | -1.44816 |
| PA2672 | -1.23836 |
| xcpR | -0.28436 |
| xcpU | -0.50274 |
| xcpV | -0.19849 |
| xcpY | -0.22214 |
| xcpZ | -0.21626 |
| tadB | -0.16973 |
| tadD | -0.25359 |
| algR | -0.35921 |
| algL | 0.640009 |
| pslF | -1.03452 |
| PA1839 | 2.200892 |
| PA0419 | -0.92439 |
| PA0017 | -0.33728 |
| PA3680 | -0.34165 |

**Table 2.** List of target DEGs and its log fold change of expression in tobramycin treated *P. aeruginosa* biofilm (GSE9989)

| Genes | Log Fc |
| --- | --- |
| mucA | 4.094536 |
| mucB | 1.155318 |
| lexA | 3.42962 |
| gidB | -3.31909 |
| pscQ | -2.35821 |
| pscP | -3.64432 |
| pscR | -3.11905 |
| pscG | -1.75324 |
| pscH | -1.52118 |
| pscE | -3.84799 |
| pscF | -1.3676 |
| pscI | -2.27261 |
| pscJ | -1.43567 |
| pscK | -1.6767 |
| xcpQ | -2.31778 |
| xcpS | -1.40397 |
| xcpT | -0.72049 |
| xcpU | -1.33575 |
| xcpV | -3.11543 |
| xcpW | -0.69783 |
| xcpX | -0.42221 |
| xcpY | -0.5616 |

### 3.4. Gene Ontology

Gene Ontology hits of DEGs in the functional classification system of PANTHER includes: 1) Molecular functions - Catalytic activity (GO:0003824), binding (GO:0005488), transcriptional regulator (GO:0098772) and transporter activity (GO:0005215). 2) Biological process - Biological regulations (GO:0065007), cellular process (GO:0009987), localization (GO:0051179), metabolic process (GO:0008152). 3) Protein classes - Nucleic acid metabolism protein (PC00171), gene-specific transcriptional regulator (PC00264), carrier protein (PC00219), and transporter protein (PC00227).

### 3.5. Functional enrichment analysis

The DEGs enriched in the functional pathways is represented in (Figure 5). DEGs enriched in the similar pathways were clustered into groups as functional annotation clusters with significant enrichment score (Table 3). The targeted DEGs were enriched in RNA Methyltransferases, 16S rRNA 7-methylguanosine methyltransferase, alginate biosynthesis, repressors for alginate synthesis, transcriptional repressor of SOS response, type II secretary proteins, type II transport domains, translocation protein in type III secretion and type III export protein.

**Table 3:** Functional annotations of the DEGs clustered into various groups with significant enrichment score. Derived from the source DAVID Bioinformatics Resources 6.8.

| Clusters | %<br>Enriched | P-Value | Fold Enrichment | FDR |
| --- | --- | --- | --- | --- |
| **Cluster 1** |  |  |  |  |
| Bacterial secretion system | 50.00 | 9.75E-21 | 16.68 | 2.93E-20 |
| Protein secretion by type II secretion system | 30.56 | 4.38E-15 | 43.27 | 8.77E-14 |
| Protein transporter activity | 27.78 | 1.65E-14 | 52.90 | 2.65E-13 |
| Protein transport | 27.78 | 4.30E-12 | 33.91 | 1.89E-10 |
| Type II protein secretion system complex | 22.22 | 1.21E-10 | 42.90 | 1.33E-09 |
| Transport | 27.78 | 2.72E-04 | 4.31 | 0.003997 |
|  |  |  |  | Cluster 1 Enrichment Score: 12.16 |
| **Cluster 2** |  |  |  |  |
| Type II protein secretion system complex | 22.22 | 1.21E-10 | 42.90 | 1.33E-09 |
| Prokaryotic N-terminal methylation site | 13.89 | 5.95E-06 | 38.23 | 4.58E-04 |
| Methylation | 11.11 | 1.52E-04 | 35.91 | 0.003334 |
|  |  |  |  | Cluster 2 Enrichment Score: 6.32 |
| **Cluster 3** |  |  |  |  |
| Methyltransferase | 13.89 | 7.52E-04 | 11.56 | 0.007267 |
| rRNA processing | 11.11 | 9.42E-04 | 19.69 | 0.007267 |
| S-adenosyl-L-methionine | 13.89 | 9.91E-04 | 10.75 | 0.007267 |
| rRNA base methylation | 8.33 | 0.002519 | 36.88 | 0.010077 |
| Cytoplasm | 22.22 | 0.009652 | 3.16 | 0.047188 |
| Transferase | 13.89 | 0.396132 | 1.53 | 0.977778 |
|  |  |  |  | Cluster 3 Enrichment Score: 2.06 |

**Figure 5:**
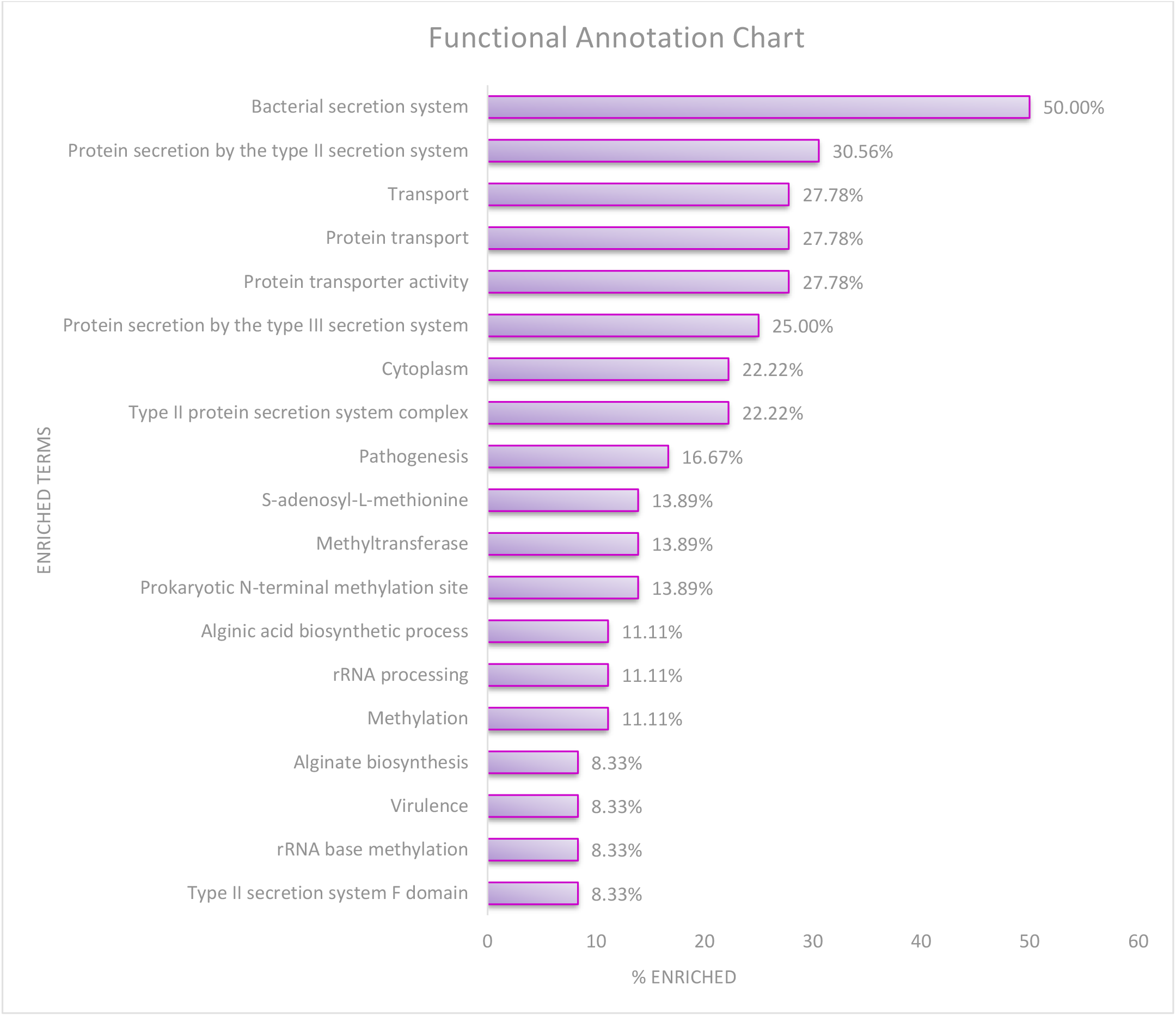
Pathway enrichments of DEGs. The functional annotations of target DEGs were performed in the DAVID Bioinformatics Resources 6.8. and represented in graph with percentage of DEGs enriched in the pathway.

## 4. Discussion

The experimental setup of *in vitro* exposure of antibiotic to *P. aeruginosa* reflects the antibiotic tolerance in chronic infections, where the response to antimicrobials diminishes due to development of adaptive resistance. Among the antibiotics evaluated for tolerance in *P. aeruginosa*, tobramycin exhibited a superior post-antibiotic effect in all MICs and was observed to be more effective by suppressing the antibiotic tolerance mechanism. Further, in silico analysis of DEGs in microarray datasets mimicking our experimental condition exposed the possible effectiveness of tobramycin in antibiotic tolerance.

In the microarray datasets, the Gene Ontological classification enabled preliminary classification of DEGs in key categories The genes of RNA Methyltransferase and methylation metabolism was enriched in catalytic activity (GO:0003824) and nucleic acid metabolism (PC00171). Regulatory genes for alginate biosynthesis pathway were observed in transcriptional regulator (GO:0098772). Other categories like transporter activity (GO:0005215), localization (GO:0051179), carrier protein (PC00219) and transporter protein (PC00227) contain genes coding for T2SS and T3SS proteins.

Among the functional enrichments observed in <u>GSE9991</u> and <u>GSE9989</u> datasets, the following enrichments play a significant role in antibiotic tolerance and virulence of *P. aeruginosa*. Methylation of 16s rRNA by Methyltransferase is a common mechanism of resistance to aminoglycosides leading to loss of affinity of the drug to the target[23]. In <u>GSE9991</u>, **RNA Methyltransferases:** PA0419, a Ribosomal RNA small subunit methyltransferase E which methylates 16s rRNA bases in 30s subunit was downregulated in the test samples. PA0017 and PA3680 genes of class B and J methyltransferase also lags expression. In <u>GSE9989</u>, **16S rRNA 7-methylguanosine methyltransferase:** gidB belongs to Methyltransferase G, which involves in methylation of 7^th^ nucleotide guanosine, confers resistance to aminoglycoside by decreasing the binding affinity to its target[24]. Low expression of gidB (LogFc -3.32) and the gene expression profile of Methyltransferases suggest low incidence of resistance development during tobramycin exposure. The following observation also extend the insights in adaptive resistance mechanism and the possibility of regulation control of RNA Methyltransferases by *P. aeruginosa* during antibiotic exposure.

In chronic infections caused by *P. aeruginosa*, biofilm formation is common during the course of infection[25], which confers additional resistance from host defenses and antibiotics[26]. Some antibiotics are involved in the up-regulation of genes that are responsible for induction of alginate production (mucopolysaccharide with an altered LPS and lipid A) which results in reduced antigen presentation to the immune system[27]. In addition, biofilms of *P. aeruginosa* also contributes to antibiotic tolerance and the regulations of several biofilm forming genes will affect the persistence of the cells in antibiotics[28]. In <u>GSE9991</u>, **Alginate Biosynthesis:** algL is a lyase precursor, that participates in catabolic activity of alginic acid, which leads to deconstruction of alginate complex[29] was highly expressed. Regulatory protein of alginate biosynthesis genes, algR [30] declined in expression. pslF was low expressed (LogFc -1.03) which is one among glycosyltransferase family, involved in extracellular polysaccharide biosynthetic pathway[31]. In <u>GSE9989</u>, **Repressors for alginate synthesis:** algU is the sigma factor for alginate biosynthesis genes. mucA codes for anti-sigma factor and mucB is a negative regulator of algU[32]. High expression of mucA (LogFc 4.09) and mucB (LogFc 1.15) may downregulates the alginate biosynthesis. The following transcriptional changes observed would possibly affect the alginate production to significant level in the presence of tobramycin treatment.

Previous studies suggests that toxin-antitoxin system mediates persister cells in antibiotics. Although, studies in E.coli shows evidence for the mechanism[33], it is not clear in *P. aeruginosa*. Some of the DEGs were linked to suppress proteins of type II secretion system and type III secretion system, which participates in virulence activity of *P. aeruginosa*. In <u>GSE9991</u>, **Type II Secretary Proteins:** PA2677, PA2672 and PA0687 engages in catalytic and transporter protein activity was downregulated. Other transporter domains xcpR, xcpU, xcpV, xcpX, xcpY, xcpZ, tadB and tadD were also low expressed[34], affecting the T2SS. In <u>GSE9989</u>, **Type II Transport Domains:** xcpQ, xcpS, xcpT, xcpU, xcpV, xcpW, xcpX and xcpY which involves in efflux of toxin from xcpR, a cytosolic domain was down-regulated affecting the export of T2SS[35]. **Translocation protein in type III secretion:** pscQ, pscP and pscR are translocation protein of type III secretion system, which translocate the toxin across the host cell cytoplasmic membrane. Downregulation of pscQ negatively impact the toxin delivery to cytosol of host cell. **Type III Export protein:** pscE, pscF, pscG, pscH, pscI, pscJ, pscK are the export proteins of type III secretion system present in cytoplasmic membrane which transfers the toxin from cytosolic domain to MS ring of basal body[35]. Low expression of all these proteins, prevents toxins to reach filament, from where it is translocated into host cell. Following transcriptomic changes suggests the suppression of toxin port systems (T2SS and T3SS), which may reduce the virulence of the organism during tobramycin treatment.

Overuse of antibiotics hiked up transcriptional regulation, favoring adaptive resistance which out-turns the fall in antibiotic activity over time[36]. This study suggests the use of tobramycin for treatment of chronic *pseudomonas* infection, as *P. aeruginosa* failed to develop adaptive resistance in tobramycin and exhibited a positive transcriptomic regulation for antibiotic response. Tobramycin is restricted for systemic use due rise of creatinine level during initial days of therapy. Intensity of nephrotoxicity between aminoglycosides is poorly understood. Recent cohort study on, nephrotoxicity suggested that, tobramycin has less comparative toxicity over gentamicin[37]. Considering the *in vivo* drug response and predisposing factors, tobramycin one among the option might enable a better treatment alternative from the current drug combinations.

Transcriptional alterations in microbes are dynamic event triggered by environmental changes, which out-turns the increase in adaptive resistance. Although, adaptive resistance involves in hike of baseline MIC of the bacteria over time, genetic resistance is a function of time. It takes several generations of the bacteria to achieve genotypic resistance. The methodology of constantly switching antibiotics through *in vitro* exposure of antibiotics would enable us in deciphering which clinical isolates would be physiologically resistant, leading to alternative aminoglycoside treatment for combating chronic infections.

## Author Contributions

All the authors had contributed for the above study in several ways.

|  | Contributor 1 | Contributor 2 | Contributor 3 | Contributor 4 |
| --- | --- | --- | --- | --- |
| Concepts | ✓ |  |  |  |
| Design | ✓ | ✓ |  |  |
| Definition of intellectual content |  |  | ✓ | ✓ |
| Literature search | ✓ | ✓ |  |  |
| Experimental studies | ✓ | ✓ |  |  |
| Data acquisition | ✓ |  |  |  |
| Data analysis | ✓ |  |  |  |
| Statistical analysis | ✓ |  |  |  |
| Manuscript preparation | ✓ | ✓ |  |  |
| Manuscript editing | ✓ | ✓ | ✓ | ✓ |
| Manuscript review |  | ✓ | ✓ | ✓ |
| Guarantor |  |  | ✓ |  |

## Ethical Statement

The study did not involve any human or animal experimentations.

## Acknowledgment

We would like to express our gratitude to Department of Microbial Biotechnology, Bharathiar University for the laboratory and technical support. We are thankful to Dr. Angayarkanni for guidance during the study period.

## References

[1] Carmeli Y, Troillet N, Eliopoulos GM, Samore MH. Emergence of antibiotic-resistant Pseudomonas aeruginosa: Comparison of risks associated with different antipseudomonal agents. Antimicrob Agents Chemother 1999;43:1379–82. https://doi.org/10.1128/aac.43.6.1379.

[2] Lynch JP, Zhanel GG, Clark NM. Emergence of Antimicrobial Resistance among Pseudomonas aeruginosa : Implications for Therapy. Semin Respir Crit Care Med 2017;38:326–45. https://doi.org/10.1055/s-0037-1602583.

[3] Bassetti M, Vena A, Croxatto A, Righi E, Guery B. How to manage Pseudomonas aeruginosa infections. Drugs Context 2018;7:1–18. https://doi.org/10.7573/dic.212527.

[4] Prayle A, Smyth AR. Aminoglycoside use in cystic fibrosis: Therapeutic strategies and toxicity. Curr Opin Pulm Med 2010;16:604–10. https://doi.org/10.1097/MCP.0b013e32833eebfd.

[5] Falagas ME, Matthaiou DK, Bliziotis IA. The role of aminoglycosides in combination with a β-lactam for the treatment of bacterial endocarditis: A meta-analysis of comparative trials. J Antimicrob Chemother 2006;57:639–47. https://doi.org/10.1093/jac/dkl044.

[6] Bodmann KF. Current guidelines for the treatment of severe pneumonia and sepsis. Chemotherapy 2005;51:227–33. https://doi.org/10.1159/000087452.

[7] Dornbusch K, Olofsson C, Holm S. Postantibiotic effect and postantibiotic sub-mic effect of dirithromycin and erythromycin against respiratory tract pathogenic bacteria. Apmis 1999;107:505–13. https://doi.org/10.1111/j.1699-0463.1999.tb01586.x.

[8] Javiya V, Ghatak S, Patel K, Patel J. Antibiotic susceptibility patterns of Pseudomonas aeruginosa at a tertiary care hospital in Gujarat, India. Indian J Pharmacol 2008;40:230–4. https://doi.org/10.4103/0253-7613.44156.

[9] Karlowsky JA, Saunders MH, Harding GAJ, Hoban DJ, Zhanel GG. In vitro characterization of aminoglycoside adaptive resistance in Pseudomonas aeruginosa. Antimicrob Agents Chemother 1996;40:1387–93. https://doi.org/10.1128/aac.40.6.1387.

[10] Gilleland LB, Gilleland HE, Gibson JA, Champlin FR. Adaptive resistance to aminoglycosides antibiotics in Pseudomonas aeruginosa. J Med Microbiol 1989;29:41–50. https://doi.org/10.1099/00222615-29-1-41.

[11] Hocquet D, Vogne C, El Garch F, Vejux A, Gotoh N, Lee A, et al. MexXy-OprM efflux pump is necessary for adaptive resistance of Pseudomonas aeruginosa to aminoglycosides. Antimicrob Agents Chemother 2003;47:1371–5. https://doi.org/10.1128/AAC.47.4.1371-1375.2003.

[12] Moradali MF, Ghods S, Rehm BHA. Pseudomonas aeruginosa lifestyle: A paradigm for adaptation, survival, and persistence. Front Cell Infect Microbiol 2017;7. https://doi.org/10.3389/fcimb.2017.00039.

[13] Sequence T, Pseudomonas A. Transcriptome Sequence of Antibiotic-Treated Pseudomonas aeruginosa. Am Soc Microbiol 2019;8:1–2. https://doi.org/10.1128/MRA.01367-18.

[14] Lonergan Z, Nairn B, Wang J, Hsu Y-P, Hesse L, Beavers W, et al. An Acinetobacter baumannii, Zinc-Regulated Peptidase Maintains Cell Wall Integrity during Immune-Mediated Nutrient Sequestration. Cell Rep 2019;26:2009-2018.e6. https://doi.org/10.1016/j.celrep.2019.01.089.

[15] Berti AD, Wergin JE, Girdaukas GG, Hetzel SJ, Sakoulas G, Rose WE. Altering the Proclivity towards Daptomycin Resistance in Methicillin-Resistant Staphylococcus aureus Using Combinations with Other Antibiotics. Antimicrob Agents Chemother 2012;56:5046 LP–5053. https://doi.org/10.1128/AAC.00502-12.

[16] A. M, J. C, T. M, U. K, I. R, P. I, et al. Evaluation of Matrix-Assisted Laser Desorption Ionization-Time-of-Flight Mass Spectrometry in Comparison to 16S rRNA Gene Sequencing for Species Identification of Nonfermenting Bacteria. J Clin Microbiol 2008;46:1946–54. https://doi.org/10.1128/JCM.00157-08.

[17] Anderson GG, Moreau-Marquis S, Stanton BA, O’Toole GA. In Vitro Analysis of Tobramycin-Treated Pseudomonas aeruginosa Biofilms on Cystic Fibrosis-Derived Airway Epithelial Cells. Infect Immun 2008;76:1423 LP–1433. https://doi.org/10.1128/IAI.01373-07.

[18] Xia J, Gill EE, Hancock REW. NetworkAnalyst for statistical, visual and network-based meta-analysis of gene expression data. Nat Protoc 2015;10:823–44. https://doi.org/10.1038/nprot.2015.052.

[19] Mi H, Thomas P. PANTHER pathway: an ontology-based pathway database coupled with data analysis tools. Methods Mol Biol 2009;563:123–40. https://doi.org/10.1007/978-1-60761-175-2_7.

[20] Huang DW, Sherman BT, Lempicki RA. Bioinformatics enrichment tools: paths toward the comprehensive functional analysis of large gene lists. Nucleic Acids Res 2009;37:1–13. https://doi.org/10.1093/nar/gkn923.

[21] Winsor GL, Griffiths EJ, Lo R, Dhillon BK, Shay JA, Brinkman FSL. Enhanced annotations and features for comparing thousands of Pseudomonas genomes in the Pseudomonas genome database. Nucleic Acids Res 2016;44:D646–53. https://doi.org/10.1093/nar/gkv1227.

[22] Romero P, Karp P. PseudoCyc, A Pathway-Genome Database for Pseudomonas aeruginosa. Microb Physiol 2003;5:230–9. https://doi.org/10.1159/000071075.

[23] Doi Y, Arakawa Y. 16S Ribosomal RNA Methylation: Emerging Resistance Mechanism against Aminoglycosides. Clin Infect Dis 2007;45:88–94. https://doi.org/10.1086/518605.

[24] Ogasawara N, Yoshikawa H. Genes and their organization in the replication origin region of the bacterial chromosome. Mol Microbiol 1992;6:629–34. https://doi.org/10.1111/j.1365-2958.1992.tb01510.x.

[25] Yu S, Wei Q, Zhao T, Guo Y, Ma LZ. A survival strategy for Pseudomonas aeruginosa that uses exopolysaccharides to sequester and store iron to stimulate psl-dependent biofilm formation. Appl Environ Microbiol 2016;82:6403–13. https://doi.org/10.1128/AEM.01307-16.

[26] Menendez A, Brett Finlay B. Defensins in the immunology of bacterial infections. Curr Opin Immunol 2007;19:385–91. https://doi.org/10.1016/j.coi.2007.06.008.

[27] Li YH, Tian X. Quorum sensing and bacterial social interactions in biofilms. Sensors 2012;12:2519–38. https://doi.org/10.3390/s120302519.

[28] Ciofu O, Tolker-Nielsen T. Tolerance and Resistance of Pseudomonas aeruginosa Biofilms to Antimicrobial Agents-How P. aeruginosa Can Escape Antibiotics. Front Microbiol 2019;10:913. https://doi.org/10.3389/fmicb.2019.00913.

[29] Schiller NL, Monday SR, Boyd CM, Keen NT, Ohman DE. Characterization of the Pseudomonas aeruginosa alginate lyase gene (algL): Cloning, sequencing, and expression in Escherichia coli. J Bacteriol 1993;175:4780–9. https://doi.org/10.1128/jb.175.15.4780-4789.1993.

[30] Deretic V, Konyecsni WM. Control of mucoidy in Pseudomonas aeruginosa: Transcriptional regulation of algR and identification of the second regulatory gene, algQ. J Bacteriol 1989;171:3680–8. https://doi.org/10.1128/jb.171.7.3680-3688.1989.

[31] Jackson KD, Starkey M, Kremer S, Parsek MR, Wozniak DJ. Identification of psl, a locus encoding a potential exopolysaccharide that is essential for Pseudomonas aeruginosa PAO1 biofilm formation. J Bacteriol 2004;186:4466–75. https://doi.org/10.1128/JB.186.14.4466-4475.2004.

[32] Qiu D, Eisinger VM, Rowen DW, Yu HD. Regulated proteolysis controls mucoid conversion in Pseudomonas aeruginosa. Proc Natl Acad Sci U S A 2007;104:8107–12. https://doi.org/10.1073/pnas.0702660104.

[33] Kim Y, Wood TK. Toxins Hha and CspD and small RNA regulator Hfq are involved in persister cell formation through MqsR in Escherichia coli. Biochem Biophys Res Commun 2010;391:209–13. https://doi.org/10.1016/j.bbrc.2009.11.033.

[34] Korotkov K V., Sandkvist M, Hol WGJ. The type II secretion system: Biogenesis, molecular architecture and mechanism. Nat Rev Microbiol 2012;10:336–51. https://doi.org/10.1038/nrmicro2762.

[35] Ma Q, Zhai Y, Schneider JC, Ramseier TM, Saier MH. Protein secretion systems of Pseudomonas aeruginosa and P. fluorescens. Biochim Biophys Acta – Biomembr 2003;1611:223–33. https://doi.org/10.1016/S0005-2736(03)00059-2.

[36] Sandoval-Motta S, Aldana M. Adaptive resistance to antibiotics in bacteria: A systems biology perspective. Wiley Interdiscip Rev Syst Biol Med 2016;8:253–67. https://doi.org/10.1002/wsbm.1335.

[37] van Maarseveen E, van Buul-Gast MC, Abdoellakhan R, Gelinck L, Neef C, Touw D. Once-daily dosed gentamicin is more nephrotoxic than once-daily dosed tobramycin in clinically infected patients. J Antimicrob Chemother 2014;69:2581–3. https://doi.org/10.1093/jac/dku175.

